# The EBI2-oxysterol axis promotes the development of intestinal lymphoid structures and colitis

**DOI:** 10.1101/424937

**Authors:** Annika Wyss, Tina Raselli, Gérard Schmelczer, Glynis Klinke, Nathan Perkins, Martin Hersberger, Marianne R. Spalinger, Kirstin Atrott, Silvia Lang, Isabelle Frey-Wagner, Michael Scharl, Andreas W. Sailer, Oliver Pabst, Gerhard Rogler, Benjamin Misselwitz

## Abstract

The gene encoding for Epstein-Barr virus-induced G-protein coupled receptor 2 (EBI2) is a risk gene for inflammatory bowel disease (IBD). Together with its oxysterol ligand 7α,25-dihydroxycholesterol, EBI2 mediates migration and differentiation of immune cells. However, the role of EBI2 in the colonic immune system remains insufficiently studied.

We found increased mRNA expression of EBI2 and oxysterol synthesizing enzymes (CH25H, CYP7B1) in the inflamed colon of patients with ulcerative colitis and mice with acute or chronic dextran sulfate sodium (DSS) colitis. Accordingly, we detected elevated extraintestinal levels of 25-hydroxylated oxysterols, including 7α,25-dihydroxycholesterol in mice with acute colonic inflammation. Knockout of EBI2 or CH25H did not affect severity of DSS colitis; however, inflammation was decreased in male EBI2^-/-^ mice in the IL-10 colitis model.

The colonic immune system comprises mucosal lymphoid structures, which accumulate upon chronic inflammation in IL-10-deficient mice and in chronic DSS colitis. However, EBI2^-/-^ mice formed significantly less colonic lymphoid structures at baseline and showed defects in inflammation-induced accumulation of lymphoid structures.

In summary, we report induction of the EBI2-7α,25-dihydroxycholesterol axis in colitis and a role of EBI2 for the accumulation of lymphoid tissue during homeostasis and inflammation. These data implicate the EBI2-7α,25-dihydroxycholesterol axis in IBD pathogenesis.

## Introduction

Inflammatory bowel diseases (IBD), with the main forms Crohn’s disease (CD) and ulcerative colitis (UC), are chronic inflammatory conditions of the human gut. The pathogenesis of IBD is incompletely understood but genetic and environmental factors were shown to contribute to disease development and progression. Genome wide association studies (GWAS) have identified more than 240 genetic regions in the human genome affecting the risk for IBD.^1, 2^ The majority of IBD-specific single nucleotide polymorphisms (SNPs) confer an increased risk for both, CD and UC.^1^ Genes identified by GWAS provide a framework for future scientific studies addressing IBD pathogenesis.

Epstein-Barr virus-induced G protein-coupled receptor 2 (EBI2, also known as GPR183), is an IBD risk gene identified by GWAS.^1^ EBI2 exerts a crucial function for the correct activation and maturation of naïve B cells in secondary lymphoid organs.^3-5^ Oxysterols are ligands for EBI2, the most potent being 7α,25-dihydroxycholesterol (7α,25-diHC).^6, 7^ 7α,25-diHC acts as a chemoattractant, directing migration of EBI2 expressing B cells, T cells and DCs.^6-9^ A 7α,25-diHC gradient in secondary lymphoid organs seems to be important for correct positioning of immune cells and a rapid and efficient antibody response.^10^ 7α,25-diHC is produced from cholesterol via two hydroxylation steps, at position 25 and 7α, by the enzymes cholesterol 25-hydroxylase (CH25H) and cytochrome P450 family 7 subfamily member B1 (CYP7B1), respectively.

Besides B cells and DCs, EBI2 is also expressed in macrophages and natural killer cells^6^, and CH25H and CYP7B1 are expressed in immune cells and many tissues including lymph nodes, lung and colon.^6, 11^ Therefore, the oxysterol-EBI2 axis might constitute a fundamental mechanism in the regulation of the immune system and tissue homeostasis.

Intestinal immune responses are orchestrated in lymphoid tissue localized directly in the intestinal tract and draining lymph nodes. Local lymphoid tissues show considerable plasticity, varying in organization and cellular composition depending on the segments of the gut and the immune status. The colonic immune system comprises two types of secondary lymphoid organs: large colonic patches (CLP), similar to Peyer’s patches (PP) in the small intestine, and smaller structures referred to as solitary intestinal lymphoid tissue (SILT).^12^ CLP and PP develop before birth, whereas SILT develop strictly postnatally. SILT comprise a continuum of lymphoid structures ranging from nascent small immature cryptopatches (CP) to mature isolated lymphoid follicles (ILF) containing B cells.

General principles of secondary lymphoid tissue formation are shared between mesenteric lymph nodes and intestinal lymphoid tissues including CLP/PP and SILT (reviewed in ^13^). Development of these lymphoid tissues includes clustering of lymphoid tissue inducer (LTi) cells, a subclass of innate lymphoid cells (ILC), and requires an intact lymphotoxin signaling pathway. Most previous studies focused on the development of lymphoid tissue in the small intestine; in contrast, formation of SILT in the colon has been studied to a lower extent and specific factors required beyond lymphotoxin remain unknown. While in the small intestine, maturation of SILT depends on CXCL13, RANKL and CCR6/CCL20, these chemokines are not essential in the colon.^12, 14^ Furthermore, while the gut microbiota stimulates small intestinal SILT, the microbiota dampens SILT formation in the colon, an effect mediated by IL-25 and IL-23.^15^ Therefore, additional molecular factors besides those cytokines seem to be required, and very recently, a role of EBI2 expressing ILCs for the development of colonic lymphoid structures has been demonstrated.^16^

In the colon, adaptation to inflammation includes formation of additional lymphoid structures. Induction of SILT upon inflammation requires the presence of microbiota and lymphotoxin signaling but was independent from the nuclear hormone receptor ROR-γt^17^, suggesting that ILCs and LTi cells are not strictly necessary. SILT were proposed to host a flexible pool of B cells for the formation of an IgA response complementing PP and CLP with a fixed B cell pool size.^18^ However, the role of SILT in colon inflammation has not been clarified.

Given the function of EBI2 in immune cell migration, we aimed at investigating the role of EBI2 and oxysterols in the pathogenesis of intestinal inflammation and the development of colonic lymphoid tissues. Our results implicate the EBI2-oxysterol axis in colonic SILT development and inflammatory responses in the colon.

## Results

### Upregulation of gene expression of EBI2 and oxysterol synthesizing enzymes in inflamed tissue of UC patients

To test for an inflammation dependent regulation of the EBI2-oxysterol axis in the gut, we analyzed results of a whole human genome microarray performed with total RNA isolated from inflamed and non-inflamed intestinal tissue from UC patients (GEO data sets: GDS3268)^19^. RNA expression levels of the oxysterol synthesizing enzymes CH25H and CYP7B1 from inflamed tissue of UC patients were significantly higher compared to non-inflamed tissue of UC patients (p<0.05 and p<0.001) and tissue of healthy controls (p<0.001 and p<0.0001; Figure 1a). A similar increase was observed for the oxysterol receptor EBI2 (p<0.001 for inflamed vs. healthy).

**Figure 1:**
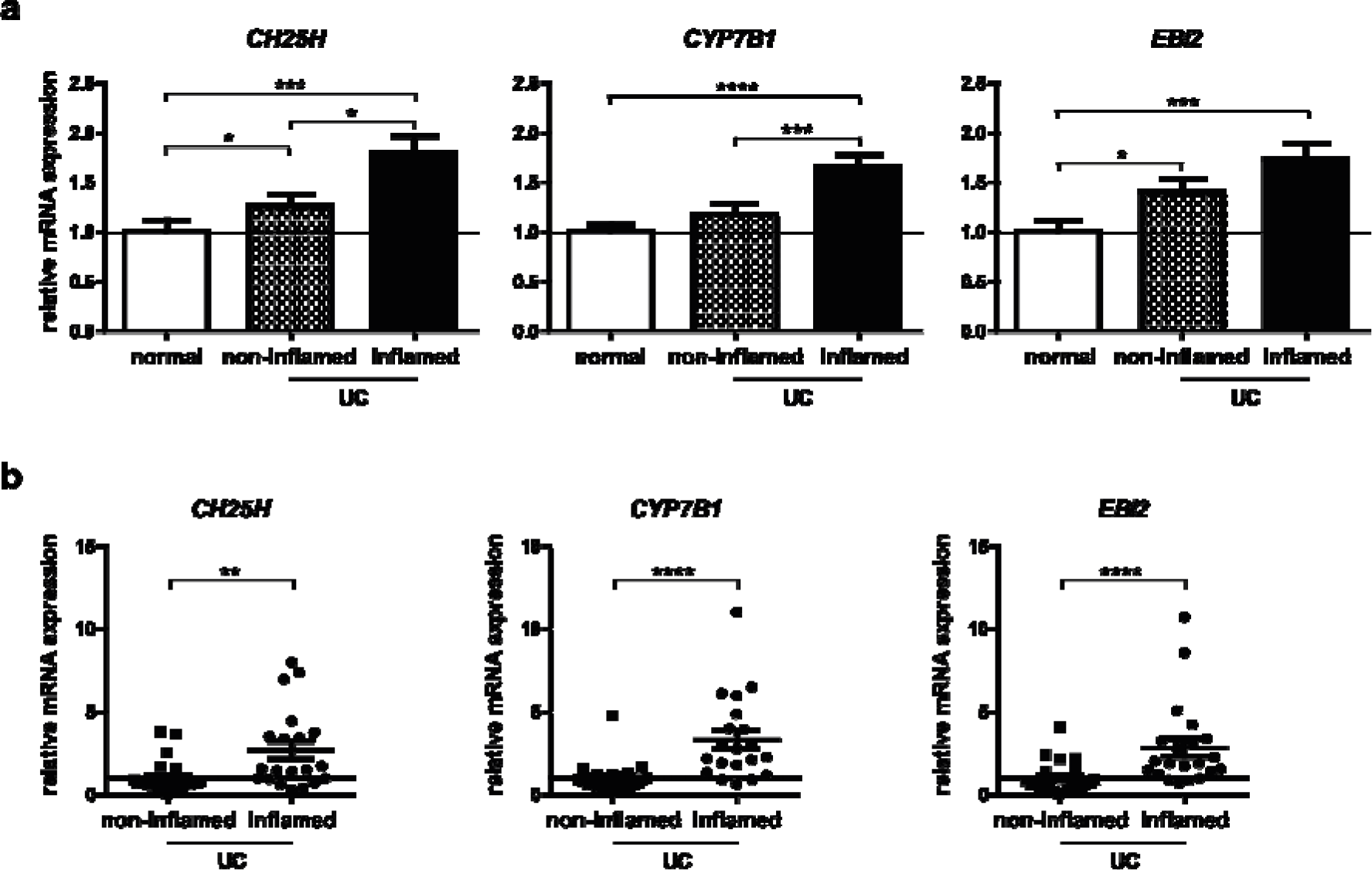
Increased expression levels of EBI2 and the oxysterol synthesizing enzymes CH25H and CYP7B1 in inflamed tissue of UC patients. **(a)** Data from a human whole genome microarray (GEO data sets: GDS3268) was analyzed regarding mRNA expression levels of *EBI2*, *CH25H* and *CYP7B1* in non-inflamed colon tissue of healthy volunteers (n=63) and non-inflamed (n=61) and inflamed (n=62) colon tissue of UC patients. The dotted line represents the mean of healthy tissue (set to 1). **(b)** mRNA expression levels from rectal biopsies of UC patients, either non-inflamed from patients with quiescent disease activity (n=24) or inflamed from patients with moderate to severe disease activity (n=20) from the Swiss IBD cohort study were determined by RT-PCR and normalized to GAPDH using the ΔΔCt method. Data shown as mean ± SEM. Statistical analysis: Mann-Whitney U test; * = p<0.05, *** = p<0.001, **** = p<0.0001.

To verify these results, we analyzed colon biopsy samples from inflamed and non-inflamed tissue of UC patients from the Swiss IBD cohort study (SIBDCS). Biopsy samples were obtained from patients with moderate to severe or quiescent UC disease activity, respectively (Supplementary Table S1).

SIBDCS samples confirmed an upregulation of *CH25H* (p<0.01), *CYP7B1* (p<0.0001) and *EBI2* (p<0.0001) mRNA levels in inflamed tissue (Figure 1b). mRNA expression levels of genes encoding pro-inflammatory cytokines (*TNF, IFNG* and *IL1B*) were used to confirm the severity of colonic inflammation (Supplementary Figure S1).

Expression levels of *CYP7B1, EBI2* and *TNF* strongly correlated (r≥0.6, p<0.001 for all correlations) and *CYP7B1* expression correlated with *CH25H* (r=0.46, p<0.05), suggesting an upregulation of the EBI2-oxysterol system in active UC in parallel with the critical cytokine *TNF* (Supplementary Table S2).

### Increased gene expression of EBI2 and oxysterol synthesizing enzymes in murine DSS colitis

To further study the role of the EBI2-oxysterol system in gut inflammation, we induced acute and chronic dextran sulfate sodium (DSS) colitis in mice. In acute colitis, expression levels of *Ebi2, Ch25h* and *Cyp7b1* were significantly increased (p<0.001; Figure 2a). This increase was more pronounced than in human samples (Figure 1), possibly reflecting the more acute and severe inflammation in DSS colitis. In chronic DSS colitis, robust up-regulation of *Ebi2* (p<0.05), *Ch25h* (p<0.05) and *Cyp7b1* (p<0.0001) was evident even though the increase was weaker in chronic than in acute inflammation, reminiscent of the human situation (Supplementary Figure S2a).

**Figure 2:**
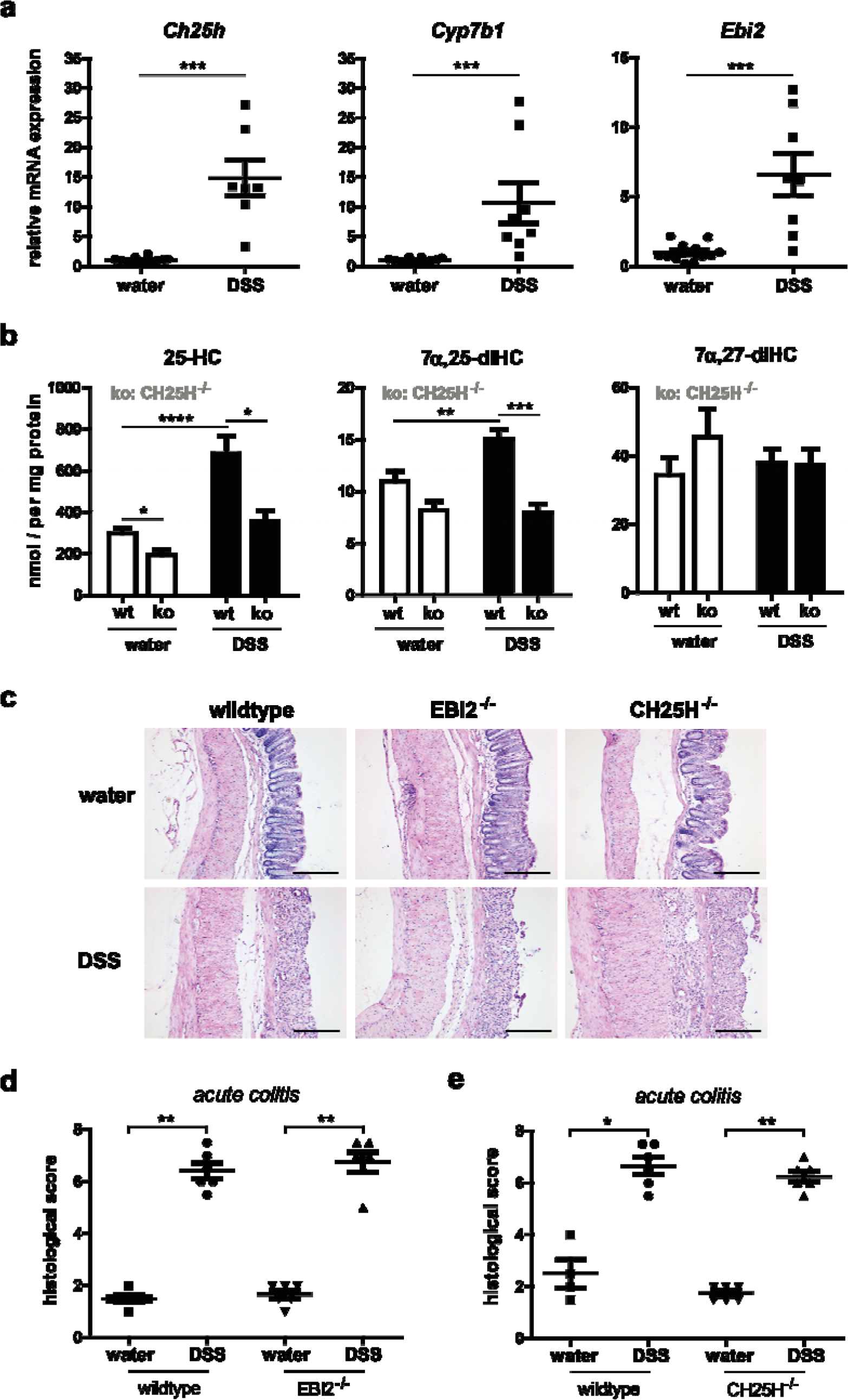
Increased expression levels of oxysterol synthesizing enzymes CH25H and CYP7B1 accompanied by elevated oxysterol levels in murine DSS colitis. Mice received DSS in drinking water for 7 days to induce acute colitis. On day 8, mice underwent colonoscopy and were sacrificed to obtain tissue samples. **(a)** mRNA expression levels of *Ch25h*, *Cyp7b1* and *Ebi2* from colon tissue of wildtype mice with acute colitis and healthy controls were determined by RT-PCR and normalized to GAPDH using the ΔΔCt method. **(b)** Oxysterol levels in liver tissue of wildtype (n=12) and CH25H^-/-^ mice (n=6) with acute colitis and healthy controls were measured by LC-MS/MS. **(c)** Histological scores for acute DSS colitis were determined on HE stained colon sections (scale bars: 200 µm) in wildtype and EBI2^-/-^ mice **(d)**, and wildtype and CH25H^-/-^ mice **(e)**. Data shown as mean ± SEM. Statistical analysis: Mann-Whitney U test; * = p<0.05, ** = p<0.01, *** = p<0.001, **** = p<0.0001.

### Increased oxysterol levels in murine colitis

To biochemically confirm effects of inflammation on CH25H and CYP7B1 activity and on oxysterol levels, we measured oxysterol concentrations of mice with acute DSS colitis and controls. For technical reasons, oxysterol levels were determined in liver and not in the gut. Eight oxysterol derivatives (hydroxycholesterol: HC; dihydroxylated cholesterols: diHC) were measured by mass spectrometry: 7α,25-diHC, 7β,25-diHC, 25-HC, 7α,27-diHC, 7β,27-diHC, 27-HC, 24S-HC and 7α,24-diHC (Figure 2b and Supplementary Figure S2b-d).

We found significantly higher levels of 25-HC and 7α,25-diHC in DSS treated mice with acute colitis compared to untreated controls (Figure 2b). Knockout of CH25H led to lower levels of 25-hydroxylated oxysterols whereas lack of EBI2 did not influence any of the oxysterols (Figure 2b and Supplementary Table S3). In a multivariate linear regression analysis (Supplementary Table S3) controlling for DSS, CH25H and EBI2 genotype, DSS treatment increased the levels of all 25-hydroxylated oxysterols (25-HC, 7α,25-diHC, 7β,25-diHC) and all 24-hydroxylated oxysterols (24S-HC, 7α,24-diHC; p≤0.0015 for all compounds). In contrast, inflammation did not seem to change 7α- or 27-hydroxylation activity.

In the same multivariate regression analysis, CH25H knockout significantly decreased all 25-hydroxylated oxysterols (7α,25-diHC, 7β,25-diHC and 25-HC, p≤0.01 for all compounds). However, as expected, the effect of CH25H knockout was not absolute and high levels of all 25-hydroxylated oxysterols remained, most likely due to the 25-hydroxylation activity of other enzymes including CYP27A1, CYP46A1 and CYP3A4.^20^

Our data thus indicate pronounced changes in liver oxysterol levels upon induction of colitis and CH25H knockout. Of note, inflammation increased levels of the EBI2 ligand 7α,25-diHC (p=0.0015) while CH25H knockout decreased its concentration (p<0.0001).

### Lack of EBI2 and CH25H does not affect severity of inflammation in the DSS colitis model

Despite the upregulation of the EBI2-oxysterol system in acute and chronic DSS colitis, knockout of neither EBI2 nor CH25H substantially decreased histological or endoscopical scoring of inflammation (Figure 2c-e, Supplementary Figure S2e-g, and data not shown). For CH25H^-/-^ mice we observed a trend towards increased inflammation indicated by significantly higher endoscopic colitis scores in acute DSS colitis (Supplementary Figure S2f) and a higher histological score in chronic DSS colitis (data not shown). Similar to wildtype animals, CH25H and EBI2 knockout mice showed increased expression levels of *Ch25h*, *Cyp7b1* and *Ebi2* in acute and chronic DSS colitis (data not shown). Taken together, in the DSS-induced colitis model, the activity of EBI2 and CH25H was not essential.

### EBI2 promotes colon inflammation in the IL-10 colitis model

Genetic defects of IL-10 or its receptor have been linked to human IBD^21^, hence we also addressed the role of EBI2 in the IL-10 model with spontaneous colitis. For this aim, we generated EBI2^-/-^IL10^-/-^mice and compared inflammatory activity to EBI2^+/+^IL10^-/-^ (subsequently named IL10^-/-^) littermate controls. In 200 days old mice, the histological colitis score of male EBI2^-/-^IL10^-/-^ mice was significantly reduced compared to IL10^-/-^ controls (p<0.01; Figure 3a, b and Supplementary Figure S3a, b). Reduced spleen weight in EBI2^-/-^IL10^-/-^ animals (p<0.05) confirmed reduced systemic inflammation in EBI2^-/-^IL10^-/-^ males (Figure 3c). Effects of EBI2 on colon inflammation were restricted to male animals, in females no EBI2 dependent changes were detected (Supplementary Figure S3c-e).

**Figure 3:**
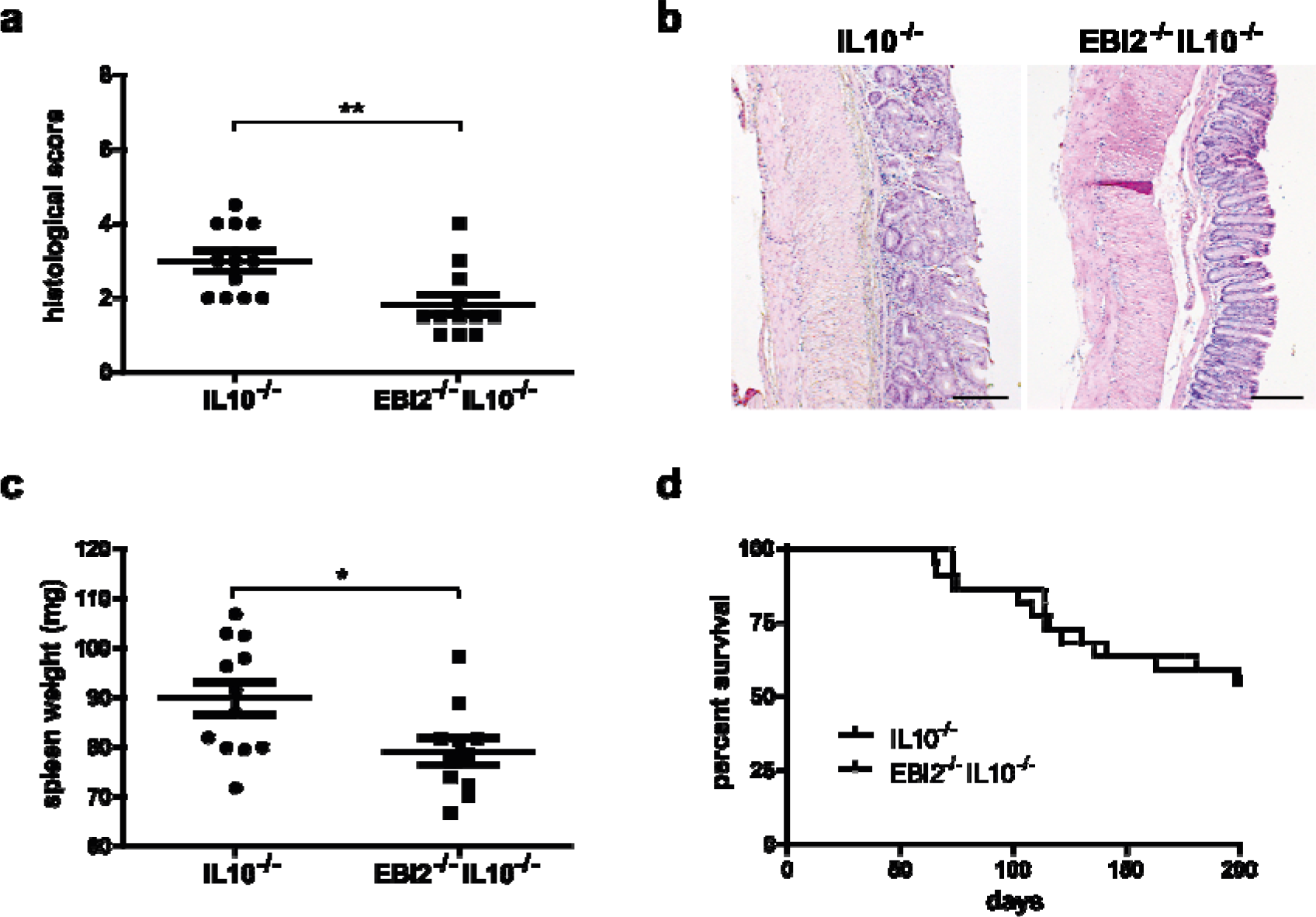
EBI2 promotes inflammation in the IL-10 colitis model. EBI2^-/-^IL10^-/-^ and IL10^-/-^ male mice were sacrificed after onset of rectal prolapse or at the age of 200 days. Histological scoring **(a)** and representative images **(b)** from HE stained colon sections from 200 days old mice. Scale bars: 200 µm. **(c)** Spleen weight of animals from (a). **(d)** Onset of prolapse in EBI2^-/-^IL10^-/-^ and IL10^-/-^ mice depicted in a survival curve. Data shown as mean ± SEM. Statistical analysis: Mann-Whitney U test; * = p<0.05, ** = p<0.01.

A minority of IL10^-/-^ animals developed rectal prolapses due to colonic inflammation; however, time to develop prolapse did not differ between EBI2^-/-^IL10^-/-^ and IL10^-/-^ in male and female animals (Figure 3d, Supplementary Figure S3e). The genotype also did not affect colon inflammation of animals with prolapse (Supplementary Figure S3a).

Testing expression levels of a panel of immune regulatory genes in 200 days old male EBI2^-/-^IL10^-/-^and IL10^-/-^ mice revealed similar expression of most cytokines and T cell regulatory genes except Tbx21 for which expression was significantly higher in EBI2^-/-^IL10^-/-^ mice (Supplementary Figure S4).

### EBI2 is required for a normal number of colonic SILT

Since EBI2 affects the distribution of immune cells in secondary lymphatic organs, we speculated that EBI2 knockout might alter the distribution of immune cells in the colon. For quantification of lymphoid structures in whole colons, B cells were visualized by B220 staining in a whole mount approach. The number of B220^+^ structures was strongly reduced in EBI2^-/-^ mice in comparison to wildtype littermate controls (p<0.05, Figure 4a, b). This difference was much stronger than experimental variation (i.e. effects of cages). Stratifying B220^+^ structures by size revealed a strongly reduced number of small and intermediate B220^+^ structures (<20’000 μm^2^: p=0.004; <100’000 μm^2^: p<0.0001) while the number of large B220^+^ structures (>100’000 μm^2^) remained unchanged (Figure 4c). Furthermore, classification into multifollicular or single structures revealed lower numbers for both types in EBI2^-/-^ mice, but with a much stronger reduction for single structures, likely representing isolated lymphoid follicles (ILF) belonging to the group of SILT (Figure 4d).

**Figure 4:**
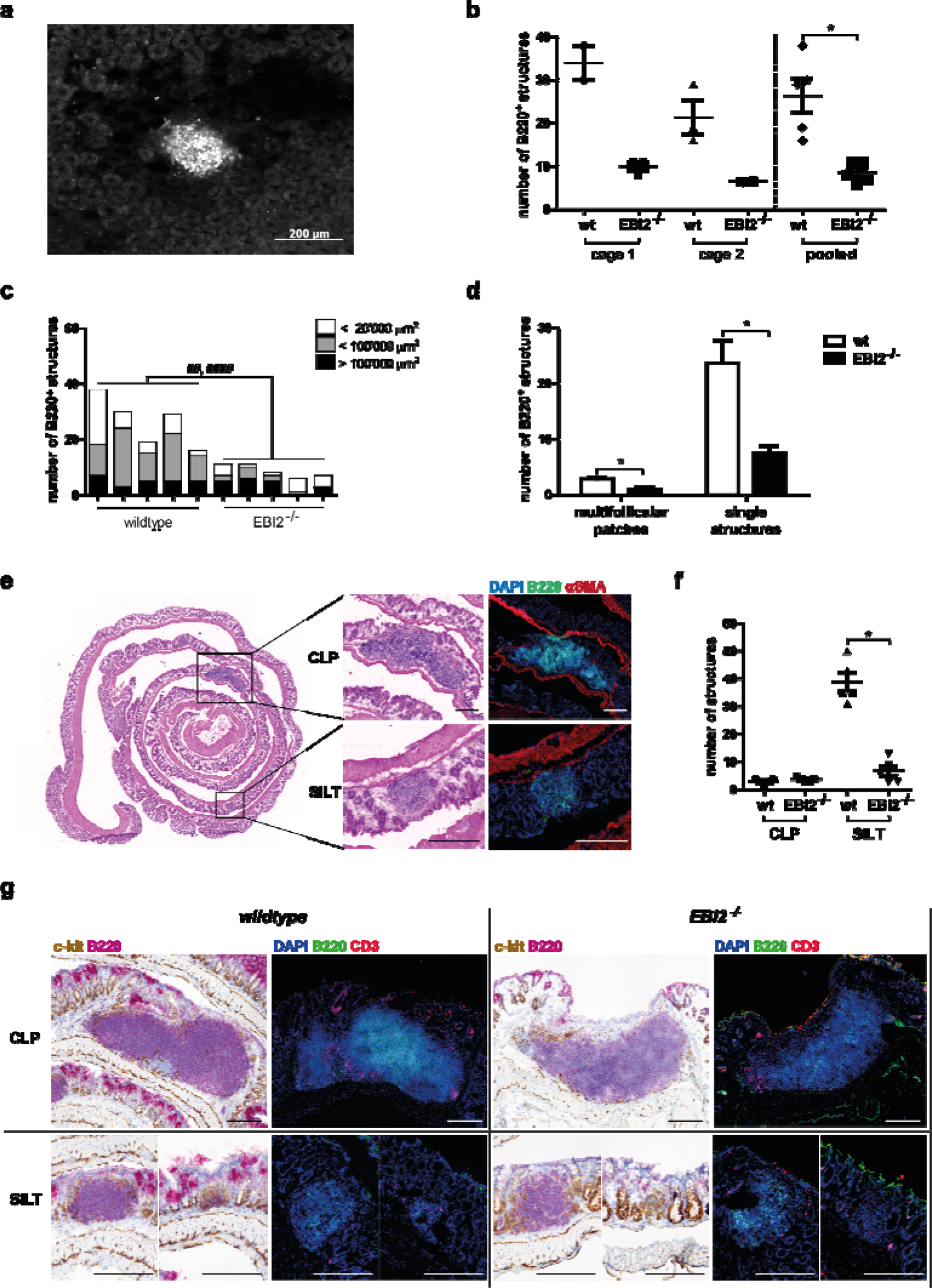
Lack of EBI2 leads to a lower number of lymphoid structures in the colon. Colonic lymphoid structures of 12 weeks old female EBI2^-/-^ and wildtype littermate mice were assessed using complementary approaches. (a) B220 B cell staining in a whole mounted colon (B220: white). (b) Quantification of B cell follicles in B220-stained whole mounted colons. (c) Stratification of B220^+^ structures by size (area). ## = p<0.01 and #### = p<0.0001 comparing structures <20’000 µm^2^ and <100’000 µm^2^ respectively with a multivariate Poisson regression using model-based t-tests. (d) Classification of B220^+^ structures by presence or absence of multiple follicles per structure. (e) HE stained colon Swiss rolls (left) and representative SILT and CLP structures stained with HE (middle) and B220 and αSMA (right); scale bars: 200 µm. (f) Lymphoid structures were quantified and categorized according to their location in 20 HE stained colon Swiss roll sections per mouse. (g) Cellular characterization of CLP and SILT with immunohistochemical staining for c-kit (brown) and B220 (pink; please note unspecific staining of alkaline phosphatase at the top of crypts) and with immunofluorescent staining for B220 (green) and CD3 (red). Scale bars: 200 µm. Data shown as mean ± SEM. Statistical analysis: Mann-Whitney U test; * = p<0.05.

Lymphoid structures can also be identified on HE stained colon “Swiss rolls”, where SILT are clearly distinguishable from CLP: SILT locate in the lamina propria, CLP between the two muscular layers and the muscularis mucosae^12^ (Figure 4e). This approach also allows for the visualization of small lymphoid structures including B cell negative cryptopatches. While the number of SILT in EBI2^-/-^ animals was reduced (p<0.05), the number of large CLP remained unchanged (Figure 4f).

Immunohistochemical stainings of the colon revealed normal structures of both SILT and CLP in EBI2^-/-^ animals with a normal distribution of B and T cells and c-kit^+^ LTi cells (Figure 4g). Taken together, our data suggest that EBI2 is required for initiation but not maturation of SILT since the few detected lymphoid structures in EBI2^-/-^ animals were indistinguishable from lymphoid structures in wildtypes.

Development of intestinal lymphoid structures is a complex process involving ROR-γt expressing ILCs, cytokines and chemokines. However, colonic expression of RORC and chemokines important for lympho-organogenesis and lymphocyte recruitment including CCL20, CXCL13 and CCL19 were not altered in EBI2^-/-^ compared to wildtype mice (Supplementary Figure S5a) suggesting that the effect of EBI2 is direct and not mediated via altered expression of any of the tested chemokines.

### EBI2 deficiency does not affect levels of B cells, IgA and microbiota composition

To further assess effects of EBI2 knockout on the colonic immune system, we quantified B and T cells by FACS. Overall, the fraction of B and T cells in the colon, mesenteric lymph nodes and spleen of EBI2^-/-^ animals was similar to wildtypes (Supplementary Figure S5b). Similarly, the substantial reduction of SILT numbers in EBI2^-/-^ mice did not significantly affect fecal IgA levels (Supplementary Figure S5c). Furthermore, overall microbiota composition of wildtype and EBI2^-/-^ animals was indistinguishable, and the microbiota of EBI2^-/-^ or wildtype animals resembled more strongly the microbiota of wildtype littermates from the same cage than animals of the same genotype housed in a different cage (data not shown). Finally, the fraction of colonic bacteria covered by IgA was similar in wildtype and EBI2^-/-^ animals (Supplementary Figure S5d). Overall, these experiments indicate an intact intestinal adaptive immune system and microbiota composition in the non-inflamed colon of EBI2^-/-^ animals.

### EBI2 promotes an increase of the number of colonic lymphoid structures in intestinal inflammation

To test for effects of EBI2 on the formation of lymphoid structures during inflammation we quantified lymphoid structures in mice with chronic DSS colitis. As expected, in wildtype mice with DSS colitis, the number of colonic lymphoid structures was approximately 2-fold higher compared to control animals (p<0.05; Figure 5a, 5b). Furthermore, in EBI2^-/-^ animals the number of lymphoid structures did not significantly increase upon DSS treatment (p=0.07, Figure 5a) and remained well below levels observed in inflamed wildtype colons even though severity of inflammation was comparable between wildtype and EBI2^-/-^ mice (Figure 2d, e). In CH25H^-/-^ mice the number of lymphoid structures at baseline was also lower compared to wildtype animals (p<0.05; Figure 5b). However, DSS-induced colonic inflammation increased the number of lymphoid structures in CH25H^-/-^ mice almost to wildtype levels. Therefore, CH25H activity seems necessary for normal development of lymphoid structures at baseline but not for an increase in chronic inflammation, indicating that other enzymes with 25-hydroxylation activity might replace CH25H. No increase in lymphoid structures was observed in acute DSS colitis (Supplementary Figure S6a, b).

**Figure 5:**
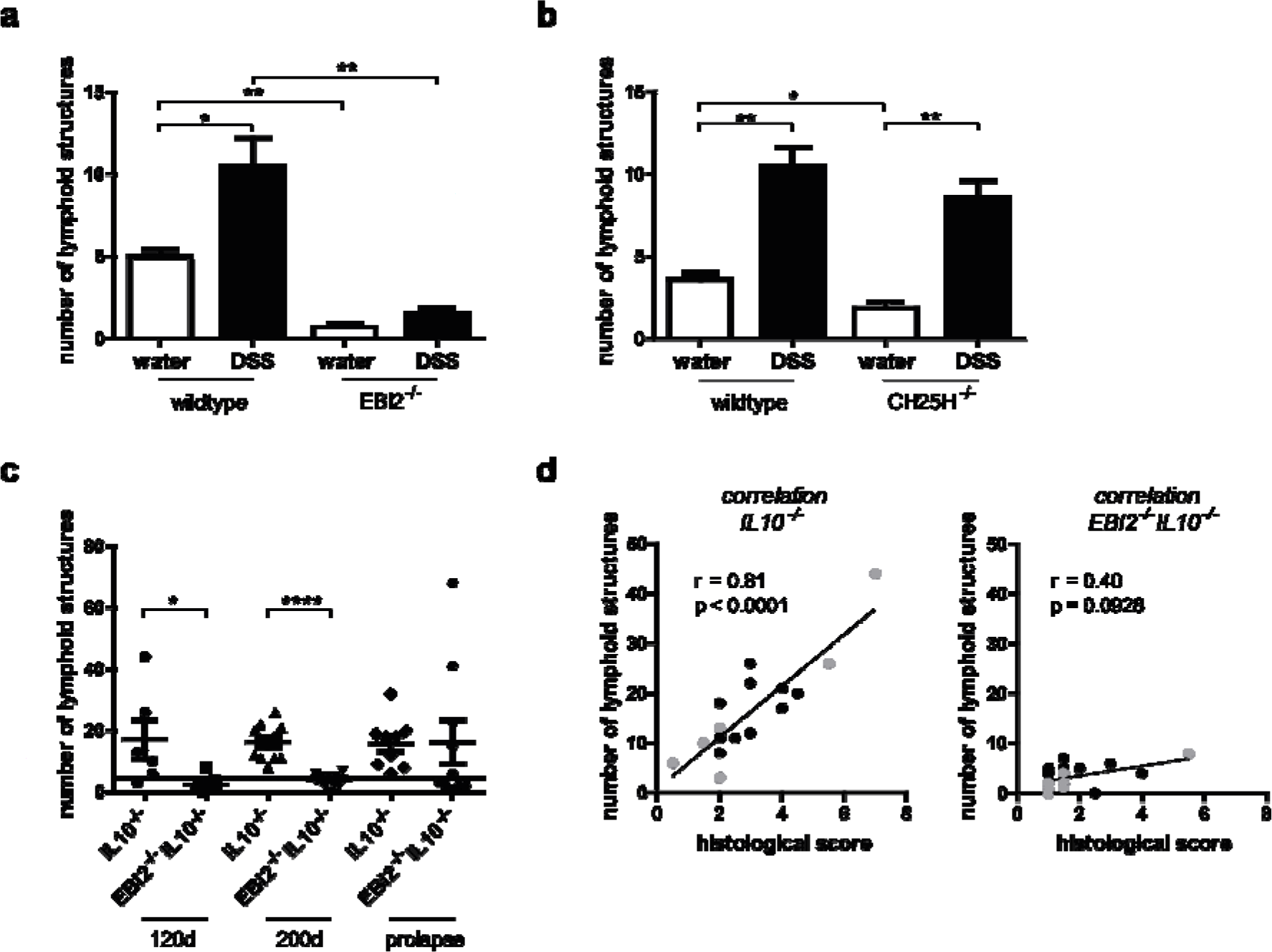
EBI2 promotes an increase in the number of colonic lymphoid structures in chronic intestinal inflammation. Lymphoid structures were quantified in representative HE stained colon sections. (a) Quantification of lymphoid structures in female wildtype and EBI2^-/-^ mice with chronic DSS colitis. (b) As in (a), but with wildtype and CH25H^-/-^ mice. (c) Quantification of lymphoid structures in EBI2^-/-^IL10^-/-^ and IL10^-/-^ male mice at the indicated ages or after occurrence of prolapse. (d) Correlation of the numbers of lymphoid structures with histological scoring of 200 days old EBI2^-/-^IL10^-/-^ and IL10^-/-^ male mice. Data shown as mean ± SEM. Statistical analysis: Mann-Whitney U test; * = p<0.05, ** = p<0.01, **** = p<0.0001; correlation analysis: Spearman R.

Compared to healthy wildtypes, the number of lymphoid structures was increased in IL10^-/-^ mice (>3-fold; Figure 5c and Supplementary Figure S7a), and the number of lymphoid structures strongly correlated with the level of intestinal inflammation in IL10^-/-^ animals (Figure 5d). However, this was not the case in EBI2^-/-^IL10^-/-^ mice, which had clearly decreased numbers of colonic lymphoid structures compared to IL10^-/-^ mice (p<0.05 and p<0.0001, at day 120 and 200, respectively). Furthermore, inflammation did not significantly increase the number of lymphoid structures in EBI2^-/-^IL10^-/-^ animals, suggesting that EBI2 is required for efficient accumulation of lymphoid structures during inflammation (Figure 5c, d and Supplementary Table S4).

While in IL10^-/-^ mice accumulation of lymphoid structures was clearly accompanied by an increase in IgA levels in fecal colon extracts, this effect was much lower in EBI2^-/-^ animals (Supplementary Figure S7b-c). Taken together, our data indicate an essential role of EBI2 in the accumulation of lymphoid structures upon colonic inflammation.

## Discussion

This study assessed the role of EBI2 and oxysterols for the development of colonic lymphoid tissue and human and murine colitis. Key observations of our study include: i) The EBI2-7α,25-diHC axis is upregulated in colitis, indicated by mRNA measurements of EBI2 and oxysterol synthesizing enzymes in human and murine colon samples and oxysterol measurements in mouse liver. ii) EBI2 is required for efficient formation of solitary intestinal lymphoid tissue (SILT) in the mouse colon. iii) EBI2 is required for accumulation of lymphoid tissue in chronic mouse colitis. iv) EBI2 increases the severity of colitis in the IL-10 colitis model but not in acute or chronic DSS colitis.

Our data demonstrate an increased mRNA expression of *EBI2*, *CH25H* and *CYP7B1* in human inflamed UC samples from two independent large data sets. Thereby, expression of *EBI2* and *CYP7B1* in the inflamed colon strongly correlated with the IBD hallmark cytokine *TNF*. These results were confirmed in murine colitis models.

Inflammation increased levels of all 25-hydroxylated oxysterols tested, including the EBI2 ligand 7α,25-diHC. This indicates that increased mRNA expression of Ch25h results in higher 25-HC production in inflammation in line with previous studies.^11, 22-24^ As expected, knockout of CH25H reduced concentrations of 25-hydroxylated oxysterols. However, concentrations of all 25-hydroxylated oxysterols remained high, in line with 25-hydroxylation activity of other enzymes such as CYP27A1, CYP46A1 and CYP3A4.^20^ Of note, for technical reasons oxysterol levels were measured in liver and not in colon tissue; any changes in oxysterol concentrations might therefore reflect changes in intestinal and/ or hepatic oxysterol synthesis due to transport of oxysterols or immune stimulants via the portal vein. Further, levels of 24-hydroxylated cholesterols, major oxysterols of the brain^25, 26^ were elevated, possibly reflecting reduced degradation in systemic inflammation. Inflammation did not affect concentrations of most 7α-hydroxylated oxysterols, even though *Cyp7b1* mRNA levels were induced 10-fold. However, local oxysterol concentrations might be much higher in the colon.

Knockout of the 7α,25-diHC receptor EBI2 decreased the number of colonic lymphoid structures, as demonstrated very recently.^16^ This reduction was shown by two independent methods: quantitatively by B220 (B cell) staining in a whole mount approach and semi-quantitatively by evaluation of consecutive HE stained sections of 100 μm distance. The latter approach allows for enumeration of cryptopatches (i.e. immature SILT lacking B cells) and a clear distinction of SILT from CLP, localized exclusively in the submucosa. Both approaches demonstrated an approximately 5-fold reduction of SILT upon EBI2 knockout. The number of SILT was also reduced in CH25H^-/-^ animals, independently confirming a role of the EBI2-7α,25-diHC axis in SILT formation.

SILT in EBI2^-/-^ and wildtype animals were morphologically indistinguishable upon staining of B, T and LTi cells, suggesting that EBI2 promotes induction but not subsequent maturation of SILT. Further, our results indicate an intact IgA response with intact bacterial IgA coating in homeostasis in EBI2^-/-^mice with a reduced number of SILTs.

Our results agree with the recent paper by Emgård et al., describing EBI2-dependent lymphoid tissue formation during homeostasis.^16^ Emgård et al. showed EBI2 expression by ILC with a LTi phenotype. LTi cells migrate towards a 7α,25-diHC gradient *in vitro* and EBI2 knockout reduced the number of cryptopatches and ILF (i.e. immature and mature SILT). Emgård et al. also demonstrated expression of CH25H and CYP7B1 by CD34^-^Podoplanin^+^ fibroblasts as a likely source for 7α,25-diHC production which would attract EBI2^+^ LTi-cells.^16^

Chronic inflammation increases the number of intestinal lymphoid tissues in the DSS colitis model.^17, 27^ Our study shows that accumulation of lymphoid structures in chronic DSS colitis strongly depends on EBI2 activity. Similarly, in IL-10 colitis, the level of inflammation increased the number of colonic lymphoid structures and EBI2 knockout abolished accumulation of SILT in inflammation (Supplementary Table S4). In contrast, CH25H knockout did not significantly affect the number of lymphoid structures in inflammation, suggesting that 7α,25-diHC concentrations are not limiting for immune cell recruitment. Several requirements for induction of SILT in inflammation, including a role of lymphotoxin, IL-22, IL-23 and CXCL13 have been identified.^17, 28, 29^ Our study describes EBI2 as a further molecule promoting the accumulation of SILT in chronic inflammation.

Even though SILT uniformly accompany colonic inflammation, it is unclear whether SILT accumulation promotes colon inflammation or whether it is a reaction to chronic inflammation, potentially enabling downregulation of inflammation or induction of tolerance.^30^ In agreement with an active role, in UC patients, lymphoid structures were found in colon specimens and larger aggregates indicated more severe inflammation.^31^ In CD, lymphoid follicles were uniformly found below fissures and aphtous lesions.^32^ Further, lymphoid follicles were associated with lymphangiectasia and perilymphangitis^33^, possibly propagating inflammation.^34^ Lymphoid follicles were also associated with high endothelial venules, facilitating homing of immune cells.^35^

Some experimental evidence also argues for pro-inflammatory effects of SILT in murine colitis: ROR-γt deficient mice displayed more colonic SILT and more severe DSS colitis compared to wildtype. Vice versa, reduction of the number of SILT by lymphotoxin neutralization decreased colitis severity.^17^ Asimilar correlation between intestinal lymphoid tissue accumulation and severity of inflammation was observed in the TNF^ΔARE^ model. Interference by anti-CCR7 treatment further increased lymphoid tissue formation and severity of inflammation.^36^ However, in our experiments, severity of chronic DSS colitis was identical in wildtype and EBI2^-/-^ mice, arguing against a pro-inflammatory role of SILT in this model.

In contrast to DSS colitis, EBI2 knockout reduces inflammation in IL-10 colitis. For unclear reasons, EBI2 deficiency only reduced inflammation in male mice. Interestingly, in a recent study with the murine TNF^ΔARE^ model, protection from ileitis was also observed in male, but not female mice with a specific pathogen free flora.^37^ Differential effects of EBI2 knockout with reduction of inflammation in IL-10 colitis but not in acute or chronic DSS colitis likely reflect different mechanisms of inflammation.^38^ In DSS colitis, pathogenesis comprises destruction of the epithelial barrier and primarily innate immune system activation. In contrast, IL-10 colitis results from lack of immune-modulatory effects of IL-10 on several immune cells.^39^ To test whether a reduced number of SILT in EBI2^-/-^IL10^-/-^ mice would reduce recruitment of pro-inflammatory T cells, we analyzed mRNA levels of several T cell markers but no difference was detected (Supplementary Figure S4).

ILCs were shown to have high EBI2 expression^16^ and ILCs lacking EBI2 failed to localize into colonic lymphoid structures.^16^ Impaired ILC recruitment might explain reduced inflammation in EBI2^-/-^IL10^-/-^ animals. The role of ILCs in IL-10 colitis has not been formally addressed; however, in a related infectious colitis model with *Campylobacter jejuni* infecting IL10^-/-^ mice, colonic ILCs were increased and depletion of ILCs abrogated colitis.^40^ ILCs have also been implicated in other models of colitis.^41-43^ In CD40 colitis, ILCs within colonic cryptopatches increased their motility shortly after induction of inflammation, resulting in ILC accumulation at the tip of the villus.^44^ The signal driving ILC movements is unclear but 7α,25-diHC produced immediately after onset of inflammation might stimulate EBI2-dependent ILC motility.

Emgård et al. also demonstrated reduced colon inflammation upon EBI2 knockout: In CD40 colitis, EBI2 expressing ILCs and myeloid cells accumulated in inflammatory foci in the colon mucosa and colitis severity was much lower in EBI2^-/-^ animals.^16^ These data, together with our data, suggest that the dependence of inflammation on EBI2 varies regarding the colitis model. EBI2 dependence would be expected for models with a critical role of ILCs including CD40 colitis^16, 42^, TRUC colitis(spontaneous colitis in RAG2^-/-^TBX21^-/-^ animals)^43^ and some infectious colitis models.^41^ In contrast, DSS colitis with breakdown of the physical barrier of the colon seems to develop independently of EBI2.

ILCs have also been implicated in human IBD since proinflammatory ILCs were found in intestinal samples.^45^ In addition, a SNP associated with the EBI2 gene increased the risk for both, UC and CD.^1^

In summary, we describe increased oxysterol synthesis in colon inflammation and a role of the EBI2-7α,25-diHC axis for generation of colonic SILT in steady state and inflammation. Our results provide further insights to SILT development in the colon, which has been substantially less studied than the small intestine. We also report a role of EBI2 in IL-10 colitis, pointing to EBI2 as a potential drug target in IBD since specific EBI2 inhibitors are available.^46^

## Materials and methods

### Human samples

Intestinal biopsies from IBD patients were obtained from the Swiss IBD Cohort Study (SIBDCS), a large, prospective nation-wide registry, started in 2007. SIBDCS is funded by the Swiss National Science Foundation and has been approved by the local ethics committee of each participating center (EK-1316). The cohort goals and methodology are described elsewhere.^47^ All patients provided written informed consent prior to inclusion into SIBDCS. Analysis of samples for the current study was approved by the scientific board of SIBDCS. Data from a whole human genome oligo microarray (GEO data sets: GDS3268)^19^ was used as a complementary data set.

### Animals

All mice were kept in a specific pathogen-free (SPF) facility in individually ventilated cages. EBI2^-/-^ mice (C57BL/6 x C129) were originally purchased from Deltagen (San Mateo, USA) and subsequently back-crossed to C57BL/6 for more than 10 generations.^6^ CH25H-deficient mice (C57/BL6) were provided by David. W. Russell, University of Texas Southwestern.^6^ In our facility, they were crossed with wildtype C57BL/6 mice and heterozygous offspring was mated to generate knockout and wildtype littermates. All animal experiments were conducted according to Swiss animal well fare law and approved by the local animal welfare authority of Zurich county (Tierversuchskommission Zürich, Zurich, Switzerland, License No. ZH256-2014).

### Colitis models

Acute dextran sulfate sodium (DSS) colitis was induced in age- and weight matched females by administration of 3% DSS (MW: 36-50 kDa; MP Biomedicals/ Thermo Fischer Scientific, Waltham, USA) in the drinking water for 7 days. Mice were sacrificed after colonoscopy on day 8. To induce chronic colitis, animals underwent 4 DSS cycles, consisting of 2% DSS administration for 7 days followed by 10 days of normal drinking water, each. Mice were sacrificed after colonoscopy 3 weeks after the last DSS cycle. Colonoscopy was performed as described previously and scored using the murine endoscopic index of colitis severity (MEICS) scoring system.^48^

IL10^-/-^ mice (C57BL/6) were crossed with EBI2^-/-^ mice; animals heterozygous for EBI2 (EBI2^+/-^IL10^-/-^) were used to generate EBI2^-/-^IL10^-/-^ and EBI2^+/+^IL10^-/-^ littermates. IL10^-/-^ animals are highly susceptible to develop spontaneous colitis. As described in previous studies^49^, 1% DSS for 4 days in drinking water at the age of 90 days was used to trigger inflammation, as the spontaneous development of colitis is dependent on the gut microbiota and is drastically reduced under SPF conditions. Mice were sacrificed at 120 or 200 days of age or upon development of rectal prolapse.

### Histological score

Colons were dissected, fixed in 4% formalin, embedded in paraffin, and cut into 5 μm sections. Deparaffinized sections were stained with hematoxylin and eosin (HE). Histological scoring for acute and chronic DSS colitis was performed as described^50^; a slightly adapted score was used for IL10^-/-^ colitis (Supplementary Table S5).

### Oxysterol measurements

#### Extraction of oxysterols from murine liver samples

Frozen liver samples from acute DSS colitis were pulverized using a CryoPrepTM CP02 (Covaris, Woburn, USA), weighed and lysed in homogenization buffer (0.9% sodium chloride) using a Qiagen TissueLyser II (Qiagen, Venlo, NL) at 4 °C. Oxysterols were extracted from the lysate by two liquid-liquid extractions using a methanol:dichloromethane (1:1) mix and a homogenization buffer:dichloromethane (1:2) mix respectively. The organic phases from both extractions were pooled and dried under nitrogen. The residue was reconstituted with ethanol containing 0.1 % formic acid and filtered before analysis on the liquid chromatography tandem-mass spectrometry (LCMS/MS) system.

#### LC-MS/MS analysis

The LC-MS/MS analysis was performed on a Dionex UltiMate 3000 RS with HPG Pump (Thermo Scientific, Waltham, USA), coupled with a Sciex Triple QuadTM 5500 mass spectrometer (AB Sciex, Zug, CH).

Detailed protocols of oxysterol extraction and LC-MS/MS analysis are provided in the supplementary methods.

### Quantification of lymphoid structures

#### Whole mount

Colons were removed intact, flushed with cold PBS, opened along the mesenteric border, and mounted, lumen facing up. Colons were then incubated two times for 10 min with warmed HBSS containing 2 mM EDTA at 37°C on a shaker to remove epithelial cells. After washing with PBS, colons were fixed with 4% Paraformaldehyde (PFA) for 1 h at 4°C. Colons were washed 5 times with TBST (1 M Tris (pH 7.2), 1 M NaCl, 0.2% Triton X-100) and blocked with TBST containing 2% rat serum for 1 h at 4°C. Colons were incubated with Cy3-conjugated anti-mouse B220 antibody (clone TIB146; provided by Oliver Pabst: Institute for Molecular Medicine, RWTH Aachen University, Aachen, Germany) in the above solution overnight at 4°C. The next day, colons were washed with TBST and mounted on glass slides. B220^+^ cell clusters were quantified under the microscope.

#### Swiss rolls

Colons were dissected and opened along the mesenteric border. Swiss rolls were prepared with the luminal side facing outwards and embedded in optimum cutting temperature (OCT) compound, and frozen in liquid nitrogen. Quantification of lymphoid structures was performed in 20 HE-stained cryosections (6 µm) per colon, approximately 100 µm apart (the protocol for immunofluorescent stainings is provided in the supplementary methods). Zeiss Axio Scan.Z1 with ZEN blue Software (Zeiss, Oberkochen, Germany) was used to scan and analyze sections. SILT and CLP were defined according to their size and localization; CLP: composed of large lymphoid follicles between the two external muscular layers and the muscularis mucosae, SILT: smaller clusters of lymphoid cells in the lamina propria.

Lymphoid structures in animals from colitis experiments were enumerated using one HE-stained section of the rolled-up colon per mouse.

If not indicated otherwise, a Zeiss Axio Imager Z2 Microscope with Axio Vision Software (Zeiss, Oberkochen, Germany) was used.

### Statistical analysis

If not indicated otherwise, Mann-Whitney U test was performed with GraphPad Prism software (GraphPad, San Diego, USA). For the multivariate linear regression analysis, appropriate modules of Matlab R2017b were used. To compare the number of B220^+^ lymphoid structures of different sizes (Figure 4c) we used R studio to generate a multivariate Poisson regression relating log(average counts) with the genotype using model-based t-tests. Data are presented as mean ± SEM. P values less than 0.05 were considered significant (*: p <0.05, **: p <0.01, ***: p <0.001, ****: p <0.0001).

## Acknowledgement

The authors would like to thank Silvia Lang for technical support, Jean-Benoît Rossel, Nicolas Fournier and Delphine Golay for help with SIBDC data and biosamples, Brian Lang for help with the multivariate linear regression analysis and the scientific committee of SIBDC for generous support.

## Author Contributions

AW, GR and BM conceived, designed, and supervised the study. AW, TR, GS, MRS, IFW, KA and SL performed experiments and/or were involved in data analysis. MH, GK and NP performed oxysterol measurements. AW and BM wrote the paper. MRS, AWS, IFW, OP, MH, MS, and GR critically revised the manuscript for important intellectual content. All authors approved the final version of the manuscript.

## Abbreviations

CD: Crohn’s disease
CH25H: Cholesterol 25-hydroxylase
CLP: Colonic patc
CP: Cryptopatch
CYP7B1: Cytochrome P450 family 7 subfamily B member 1
7a,25-diHC: 7a,25-dihydroxycholesterol
DSS: Dextran sulfate sodium
EBI2: Epstein-Barr virus-induced G protein coupled receptor 2 (GPR183)
GPR183: G protein-coupled receptor 183 (EBI2)
IBD: Inflammatory bowel disease
ILC: Innate lymphoid cell
ILF: Isolated lymphoid follicle
LC-MS/MS: Liquid chromatography tandem-mass spectrometry
LPL: Lamina propria lymhocytes
LTi: Lymphoid tissue inducer (cell)
MEICS: Murine endoscopic index of colitis severity
PP: Peyer’s patches
SILT: Solitary intestinal lymphoid tissue
SIBDCS: Swiss IBD cohort study
UC: Ulcerative colitis
25-HC: 25-hydroxycholesterol

## References

1. Jostins L, Ripke S, Weersma RK, Duerr RH, McGovern DP, Hui KY et al Host-microbe interactions have shaped the genetic architecture of inflammatory bowel disease. Nature 2012; 491(7422): 119–124.

2. de Lange KM, Moutsianas L, Lee JC, Lamb CA, Luo Y, Kennedy NA et al Genome-wide association study implicates immune activation of multiple integrin genes in inflammatory bowel disease. Nature genetics 2017; 49(2): 256–261.

3. Gatto D, Paus D, Basten A, Mackay CR, Brink R. Guidance of B cells by the orphan G protein-coupled receptor EBI2 shapes humoral immune responses. Immunity 2009; 31(2): 259–269.

4. Gatto D, Wood K, Brink R. EBI2 operates independently of but in cooperation with CXCR5 and CCR7 to direct B cell migration and organization in follicles and the germinal center. J Immunol 2011; 187(9): 4621–4628.

5. Pereira JP, Kelly LM, Xu Y, Cyster JG. EBI2 mediates B cell segregation between the outer and centre follicle. Nature 2009; 460(7259): 1122–1126.

6. Hannedouche S, Zhang J, Yi T, Shen W, Nguyen D, Pereira JP et al Oxysterols direct immune cell migration via EBI2. Nature 2011; 475(7357): 524–527.

7. Liu C, Yang XV, Wu J, Kuei C, Mani NS, Zhang L et al Oxysterols direct B-cell migration through EBI2. Nature 2011; 475(7357): 519–523.

8. Gatto D, Wood K, Caminschi I, Murphy-Durland D, Schofield P, Christ D et al The chemotactic receptor EBI2 regulates the homeostasis, localization and immunological function of splenic dendritic cells. Nat Immunol 2013; 14(8): 876.

9. Li J, Lu E, Yi T, Cyster JG. EBI2 augments Tfh cell fate by promoting interaction with IL-2-quenching dendritic cells. Nature 2016; 533(7601): 110–114.

10. Yi T, Wang X, Kelly LM, An J, Xu Y, Sailer AW et al Oxysterol gradient generation by lymphoid stromal cells guides activated B cell movement during humoral responses. Immunity 2012; 37(3): 535–548.

11. Bauman DR, Bitmansour AD, McDonald JG, Thompson BM, Liang G, Russell DW. 25-Hydroxycholesterol secreted by macrophages in response to Toll-like receptor activation suppresses immunoglobulin A production. Proc Natl Acad Sci U S A 2009; 106(39): 16764–16769.

12. Baptista AP, Olivier BJ, Goverse G, Greuter M, Knippenberg M, Kusser K et al Colonic patch and colonic SILT development are independent and differentially regulated events. Mucosal Immunol 2013; 6(3): 511–521.

13. Buettner M, Lochner M. Development and Function of Secondary and Tertiary Lymphoid Organs in the Small Intestine and the Colon. Front Immunol 2016; 7: 342.

14. Knoop KA, Butler BR, Kumar N, Newberry RD, Williams IR. Distinct developmental requirements for isolated lymphoid follicle formation in the small and large intestine: RANKL is essential only in the small intestine. Am J Pathol 2011; 179(4): 1861–1871.

15. Donaldson DS, Bradford BM, Artis D, Mabbott NA. Reciprocal regulation of lymphoid tissue development in the large intestine by IL-25 and IL-23. Mucosal Immunol 2015; 8(3): 582–595.

16. Emgard J, Kammoun H, Garcia-Cassani B, Chesne J, Parigi SM, Jacob JM et al Oxysterol Sensing through the Receptor GPR183 Promotes the Lymphoid-Tissue-Inducing Function of Innate Lymphoid Cells and Colonic Inflammation. Immunity 2018; 48(1): 120–132 e128.

17. Lochner M, Ohnmacht C, Presley L, Bruhns P, Si-Tahar M, Sawa S et al Microbiota-induced tertiary lymphoid tissues aggravate inflammatory disease in the absence of RORgamma t and LTi cells. J Exp Med 2011; 208(1): 125–134.

18. Knoop KA, Newberry RD. Isolated Lymphoid Follicles are Dynamic Reservoirs for the Induction of Intestinal IgA. Front Immunol 2012; 3: 84.

19. Noble CL, Abbas AR, Cornelius J, Lees CW, Ho GT, Toy K et al Regional variation in gene expression in the healthy colon is dysregulated in ulcerative colitis. Gut 2008; 57(10): 1398–1405.

20. Honda A, Miyazaki T, Ikegami T, Iwamoto J, Maeda T, Hirayama T et al Cholesterol 25-hydroxylation activity of CYP3A. J Lipid Res 2011; 52(8): 1509–1516.

21. Kotlarz D, Beier R, Murugan D, Diestelhorst J, Jensen O, Boztug K et al Loss of interleukin-10 signaling and infantile inflammatory bowel disease: implications for diagnosis and therapy. Gastroenterology 2012; 143(2): 347–355.

22. Diczfalusy U, Olofsson KE, Carlsson AM, Gong M, Golenbock DT, Rooyackers O et al Marked upregulation of cholesterol 25-hydroxylase expression by lipopolysaccharide. J Lipid Res 2009; 50(11): 2258–2264.

23. Blanc M, Hsieh WY, Robertson KA, Kropp KA, Forster T, Shui G et al The transcription factor STAT-1 couples macrophage synthesis of 25-hydroxycholesterol to the interferon antiviral response. Immunity 2013; 38(1): 106–118.

24. Xiang Y, Tang JJ, Tao W, Cao X, Song BL, Zhong J. Identification of Cholesterol 25-Hydroxylase as a Novel Host Restriction Factor and a Part of the Primary Innate Immune Responses against Hepatitis C Virus Infection. J Virol 2015; 89(13): 6805–6816.

25. Meljon A, Theofilopoulos S, Shackleton CH, Watson GL, Javitt NB, Knolker HJ et al Analysis of bioactive oxysterols in newborn mouse brain by LC/MS. J Lipid Res 2012; 53(11): 2469–2483.

26. Russell DW, Halford RW, Ramirez DM, Shah R, Kotti T. Cholesterol 24-hydroxylase: an enzyme of cholesterol turnover in the brain. Annu Rev Biochem 2009; 78: 1017–1040.

27. Okayasu I, Hatakeyama S, Yamada M, Ohkusa T, Inagaki Y, Nakaya R. A novel method in the induction of reliable experimental acute and chronic ulcerative colitis in mice. Gastroenterology 1990; 98(3): 694–702.

28. Ota N, Wong K, Valdez PA, Zheng Y, Crellin NK, Diehl L et al IL-22 bridges the lymphotoxin pathway with the maintenance of colonic lymphoid structures during infection with Citrobacter rodentium. Nat Immunol 2011; 12(10): 941–948.

29. Marchesi F, Martin AP, Thirunarayanan N, Devany E, Mayer L, Grisotto MG et al CXCL13 expression in the gut promotes accumulation of IL-22-producing lymphoid tissue-inducer cells, and formation of isolated lymphoid follicles. Mucosal Immunol 2009; 2(6): 486–494.

30. McNamee EN, Rivera-Nieves J. Ectopic Tertiary Lymphoid Tissue in Inflammatory Bowel Disease: Protective or Provocateur? Front Immunol 2016; 7: 308.

31. Yeung MM, Melgar S, Baranov V, Oberg A, Danielsson A, Hammarstrom S et al Characterisation of mucosal lymphoid aggregates in ulcerative colitis: immune cell phenotype and TcR-gammadelta expression. Gut 2000; 47(2): 215–227.

32. Fujimura Y, Kamoi R, Iida M. Pathogenesis of aphthoid ulcers in Crohn’s disease: correlative findings by magnifying colonoscopy, electron microscopy, and immunohistochemistry. Gut 1996; 38(5): 724–732.

33. Sura R, Colombel JF, Van Kruiningen HJ. Lymphatics, tertiary lymphoid organs and the granulomas of Crohn’s disease: an immunohistochemical study. Aliment Pharmacol Ther 2011; 33(8): 930–939.

34. Van Kruiningen HJ, Colombel JF. The forgotten role of lymphangitis in Crohn’s disease. Gut 2008; 57(1): 1–4.

35. Horjus Talabur Horje CS, Smids C, Meijer JW, Groenen MJ, Rijnders MK, van Lochem EG et al High endothelial venules associated with T cell subsets in the inflamed gut of newly diagnosed inflammatory bowel disease patients. Clin Exp Immunol 2017; 188(1): 163–173.

36. McNamee EN, Masterson JC, Jedlicka P, Collins CB, Williams IR, Rivera-Nieves J. Ectopic lymphoid tissue alters the chemokine gradient, increases lymphocyte retention and exacerbates murine ileitis. Gut 2013; 62(1): 53–62.

37. Schaubeck M, Clavel T, Calasan J, Lagkouvardos I, Haange SB, Jehmlich N et al Dysbiotic gut microbiota causes transmissible Crohn’s disease-like ileitis independent of failure in antimicrobial defence. Gut 2016; 65(2): 225–237.

38. Jones-Hall YL, Grisham MB. Immunopathological characterization of selected mouse models of inflammatory bowel disease: Comparison to human disease. Pathophysiology 2014; 21(4): 267–288.

39. Keubler LM, Buettner M, Hager C, Bleich A. A Multihit Model: Colitis Lessons from the Interleukin-10-deficient Mouse. Inflamm Bowel Dis 2015; 21(8): 1967–1975.

40. Malik A, Sharma D, St Charles J, Dybas LA, Mansfield LS. Contrasting immune responses mediate Campylobacter jejuni-induced colitis and autoimmunity. Mucosal Immunol 2014; 7(4): 802–817.

41. Buonocore S, Ahern PP, Uhlig HH, Ivanov, II, Littman DR, Maloy KJ et al Innate lymphoid cells drive interleukin-23-dependent innate intestinal pathology. Nature 2010; 464(7293): 1371–1375.

42. Vonarbourg C, Mortha A, Bui VL, Hernandez PP, Kiss EA, Hoyler T et al Regulated expression of nuclear receptor RORgammat confers distinct functional fates to NK cell receptor-expressing RORgammat(+) innate lymphocytes. Immunity 2010; 33(5): 736–751.

43. Powell N, Walker AW, Stolarczyk E, Canavan JB, Gokmen MR, Marks E et al The transcription factor T-bet regulates intestinal inflammation mediated by interleukin-7 receptor+ innate lymphoid cells. Immunity 2012; 37(4): 674–684.

44. Pearson C, Thornton EE, McKenzie B, Schaupp AL, Huskens N, Griseri T et al ILC3 GM-CSF production and mobilisation orchestrate acute intestinal inflammation. Elife 2016; 5: e10066.

45. Geremia A, Arancibia-Carcamo CV, Fleming MP, Rust N, Singh B, Mortensen NJ et al IL-23-responsive innate lymphoid cells are increased in inflammatory bowel disease. J Exp Med 2011; 208(6): 1127–1133.

46. Gessier F, Preuss I, Yin H, Rosenkilde MM, Laurent S, Endres R et al Identification and characterization of small molecule modulators of the Epstein-Barr virus-induced gene 2 (EBI2) receptor. J Med Chem 2014; 57(8): 3358–3368.

47. Pittet V, Juillerat P, Mottet C, Felley C, Ballabeni P, Burnand B et al Cohort profile: the Swiss Inflammatory Bowel Disease Cohort Study (SIBDCS). Int J Epidemiol 2009; 38(4): 922–931.

48. Becker C, Fantini MC, Neurath MF. High resolution colonoscopy in live mice. Nat Protoc 2006; 1(6): 2900–2904.

49. Michael S, Keubler LM, Smoczek A, Meier M, Gunzer F, Pohlmann C et al Quantitative phenotyping of inflammatory bowel disease in the IL-10-deficient mouse by use of noninvasive magnetic resonance imaging. Inflamm Bowel Dis 2013; 19(1): 185–193.

50. Hausmann M, Obermeier F, Paper DH, Balan K, Dunger N, Menzel K et al In vivo treatment with the herbal phenylethanoid acteoside ameliorates intestinal inflammation in dextran sulphate sodium-induced colitis. Clin Exp Immunol 2007; 148(2): 373–381.

